# Phenotyping of lymphoproliferative tumours generated in xenografts of non-small cell lung cancer

**DOI:** 10.1101/2023.01.24.520089

**Authors:** David R. Pearce, Ayse U. Akarca, Roel P. H. De Maeyer, Emily Kostina, Ariana Huebner, Monica Sivakumar, Takahiro Karasaki, Kavina Shah, Sam M. Janes, Nicholas McGranahan, Venkat Reddy, Arne N. Akbar, David A. Moore, Teresa Marafioti, Charles Swanton, Robert E. Hynds

**Author notes:** **Correspondence:** Robert E Hynds, Charles Swanton, Teresa Marafioti.

## Abstract

Patient-derived xenograft (PDX) models involve the engraftment of tumour tissue in immunocompromised mice and represent an important pre-clinixtcal oncology research. A limitation of non-small cell lung cancer (NSCLC) PDX model derivation in NOD-*scid* IL2Rgamma^null^ (NSG) mice is that a subset of initial engraftments are of lymphocytic, rather than tumour origin. In the lung TRACERx PDX pipeline, lymphoproliferations occurred in 17.8% of lung adenocarcinoma and 10% of lung squamous cell carcinoma transplantations, despite none of these patients having a prior or subsequent clinical history of lymphoproliferative disease. Lymphoproliferations were predominantly human CD20+ B cells and had the immunophenotype expected for post-transplantation diffuse large B cell lymphoma. All lymphoproliferations expressed Epstein-Barr-encoded RNAs (EBER). Analysis of immunoglobulin light chain gene rearrangements in three tumours where multiple tumour regions had resulted in lymphoproliferations suggested that each had independent clonal origins. Overall, these data suggest the presence of B cell clones with lymphoproliferative potential within primary NSCLC tumours that are under continuous immune surveillance. Since these cells can be expanded following transplantation into NSG mice, our data highlight the value of quality control measures to identify lymphoproliferations within xenograft pipelines and support the incorporation of strategies to minimise lymphoproliferations during the early stages of xenograft establishment pipelines. To present the histology data herein, we developed a Python-based tool for generating patient-level pathology overview figures from whole-slide image files; PATHOverview is available on GitHub (https://github.com/EpiCENTR-Lab/PATHOverview).

## INTRODUCTION

NOD-*scid* interleukin (IL) 2 receptor gamma chain null (NSG) mice are severely immunocompromised in both the innate and adaptive immune responses, with deficiency of functional B, T and NK cells, reduced dendritic cell and macrophage function, and extensive cytokine abnormalities^1^. Immunodeficiency in these mice facilitates the engraftment of human tissues and, in the context of cancer, offers the opportunity for individualised patient-derived xenograft (PDX) models of cancer progression and treatment response^2^.

As for other cancer types, only a fraction of implanted non-small cell lung cancers engraft in the murine host and proliferate to establish PDX lines^3–7^. Understanding why some tumours engraft and others do not is a priority, particularly as xenograft formation can correlate with clinical outcomes and some have advocated guiding patient treatment using PDX models. Although some tumour transplantations simply never form xenografts within the lifespan of the mouse, a significant cause of xenograft failure in PDX pipelines is the formation of non-tumour xenografts, which have been reported in studies of a wide range of cancer types, including liver^8^, breast^9^, gastrointestinal^9–12^, pancreatic^9^, bladder^9^, renal^9^, prostate^13^ and ovarian^14^ cancers. Similarly, this issue has been noted in lung cancer PDX models using a variety of immunocompromised mouse strains^15–17^.

We have recently reported the outcomes from a PDX generation pipeline initiated from multi-regional primary non-small cell lung cancer (NSCLC) tumour tissue within the lung TRACERx prospective cohort study^18^. We found that 16/145 NSCLC transplantations (from 13 of 44 patients) resulted in CD45+ xenograft formation^19^. Here, we report phenotyping of these CD45+ lymphoproliferations using histology, immunohistochemistry, PCR and flow cytometry approaches.

## RESULTS

### CD45+ lymphoproliferations in a NSCLC PDX model pipeline

PDX generation in NSG mice was performed through subcutaneous injection of multi-region, spatially independent biopsies of NSCLC tumours^19^. Tissue samples were obtained via the lung TRACERx study from patients undergoing surgical resections, with no patients having received neoadjuvant therapy^20^. Immunohistochemistry (IHC) revealed that 15 xenografts did not express keratin but did express human CD45^19^. Tissue from a further xenograft (CRUK0733 Region 1; R1) was not available but it was determined to be a lymphoproliferation due to its lack of shared mutations with the patient’s NSCLC in a targeted sequencing assay, meaning that overall 16/145 (11.0%) of transplanted tumour regions gave rise to lymphoproliferations (Figure 1A;^19^). Following multi-region xenograft establishment, three tumours gave rise to more than one lymphoproliferation from different spatial regions, two of these tumours also gave rise to a NSCLC PDX model from a further spatial tumour region.

**Figure 1:**
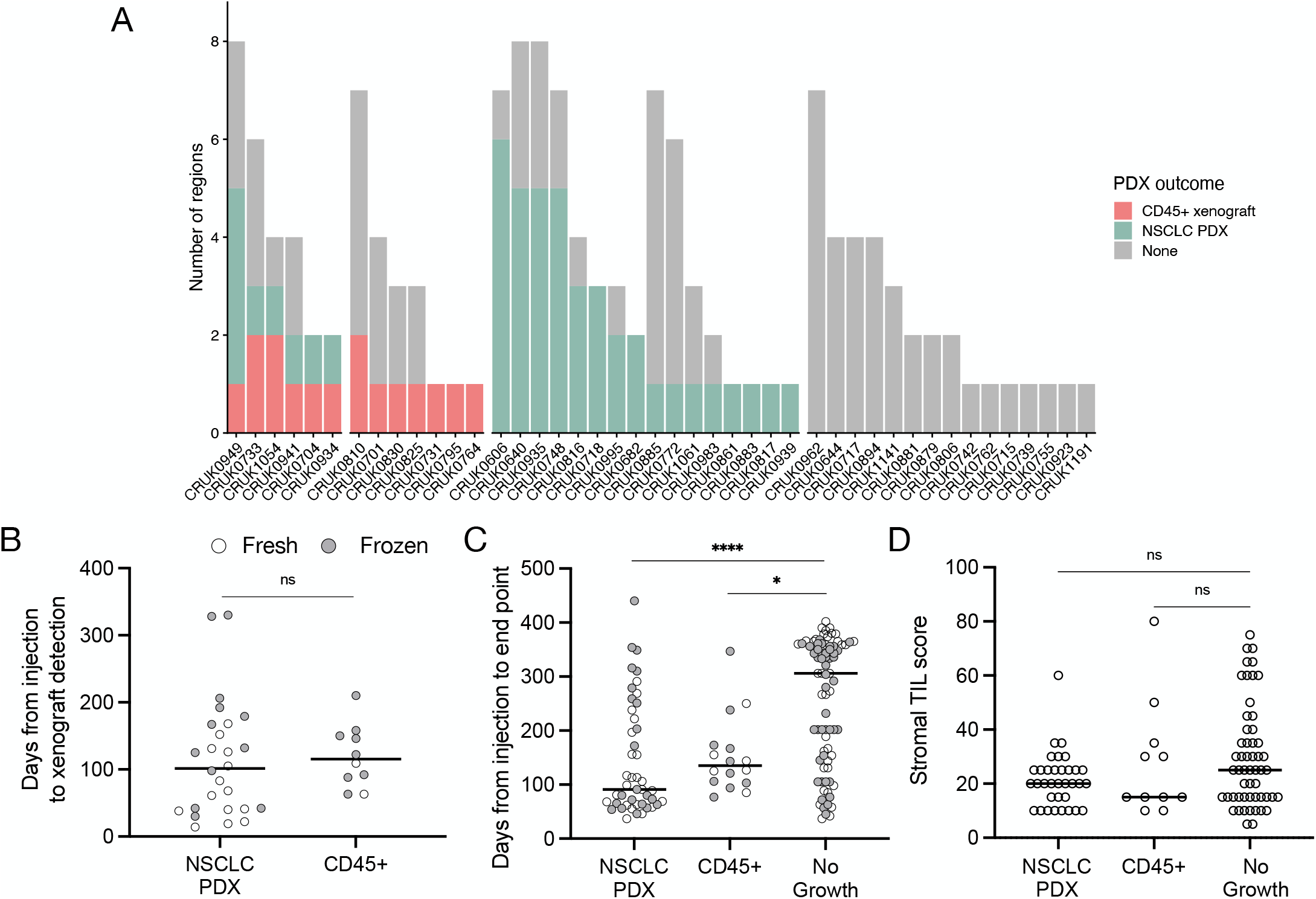
Lymphoprolifera3on forma3on in a mul3-region non-small cell lung cancer pa3ent-derived model program. A) Frequency of CD45+ xenogra6s in the lung TRACERx PDX program. B) QuanFficaFon of the Fme elapsed between injecFon and xenogra6 detecFon (ns, p = 0.33, two-tailed Mann-Whitney test). Points are coloured by the prior cryopreservaFon status of the injected tumour material. C) QuanFficaFon of the Fme elapsed between injecFon and experimental end point (xenogra6 harvest or, in the event of no xenogra6 formaFon, mouse terminaFon; ns, two-tailed Mann-Whitney test). Points are coloured by the prior cryopreservaFon status of the injected tumour material. D) Tumour infiltraFng lymphocyte score in paFent Fssue (NSCLC PDX vs no growth, p = 0.258; NSCLC PDX vs CD45+, p > 0.99; No growth vs CD45+, p > 0.99; Kruskal-Wallis test).

8/45 lung adenocarcinoma (LUAD) regions generated lymphoproliferations compared to 6/60 lung squamous cell carcinoma (LUSC) regions (ns, Chi-square test), while two lymphoproliferations arose from other NSCLC histologies (Figure 1A). We found no difference between the time taken to the emergence of palpable tumours between samples that generated lymphoproliferations versus those that generated NSCLC PDXs (Figure 1B; median 115.5 versus 101.5 days, respectively; p = 0.33, two-tailed Mann-Whitney test). Likewise, there was no difference between the time between injection of tumour material and the harvest of tumours between samples that generated lymphoproliferations versus those that generated NSCLC PDXs (Figure 1C; median 135 vs 91 days, respectively; p > 0.99, Kruskal-Wallis test). Further, 6 out of 68 (8.8%) freshly injected patient samples gave rise to lymphoproliferations compared to 10 out of 77 (13.0%) cryopreserved samples (ns, Chi-square test). Where data were available, we analysed the frequency of stromal tumour infiltrating lymphocytes in regional histology samples from our patient cohort and found no association between infiltration and lymphoproliferation (Figure 1D; ns, Kruskal-Wallis test). Clinically, none of the patients whose tumours gave rise to lymphoproliferations had a history or family history of lymphoma, nor did any of these patients relapse with lymphoproliferative disease.

### CD45+ xenografts from NSCLC are large B cell lymphomas with a post-transplant-like immunophenotype

Review of hematoxylin and eosin stained tissue sections revealed that CD45+ lymphoproliferations shared many common features independent of the tumour region or patient of origin. They contained large, atypical cells with pleomorphic nuclei containing prominent nucleoli that were mixed with mitoses and areas of necrosis. Independent xenografts exhibited variable extent of necrosis. An example patient is shown in Figure 2 and data for all patients – presented using a new Python-based tool for assembling pathology images, PATHOverview (see Methods) – are shown in Supplementary File 1.

**Figure 2:**
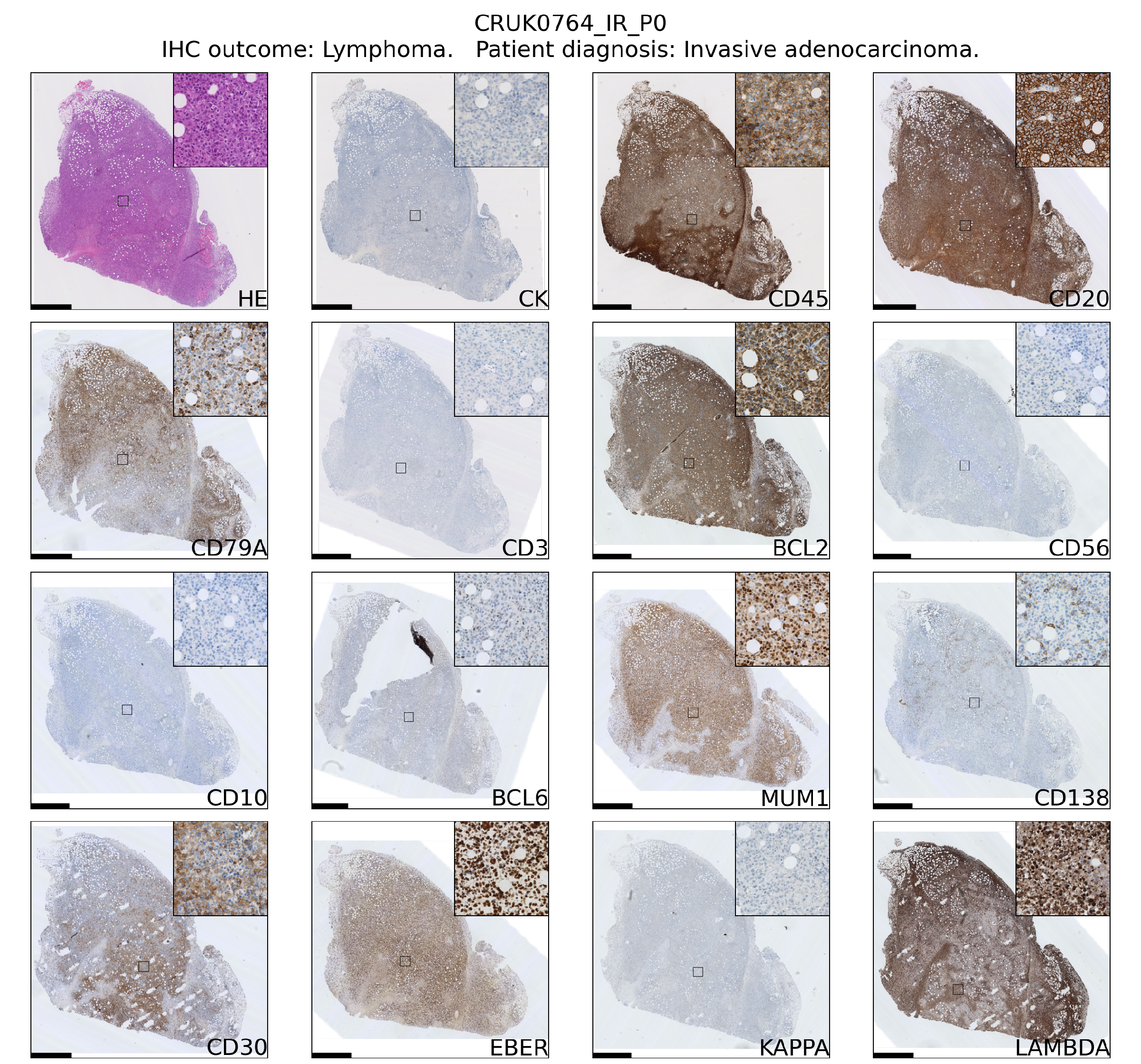
Immunophenotyping of an example case (the CRUK0764 inter-region [IR] xenograD) of lymphoprolifera3on forma3on in an NSG mouse injected subcutaneously with NSCLC. Scale bar = 1 mm. Inset image is 250 μm in width.

Conventional immunohistochemistry was carried out to characterise the immunophenotype of lymphoproliferations. Cells expressed human CD45, CD20 and CD79a, indicating their B cell identity, and in all cases were positive for Bcl-2 and CD30. They were largely negative for CD56 and the germinal centre-associated markers CD10 and Bcl-6^21^, although occasionally Bcl-6 expression was observed in a subset of the atypical cells. MUM1, a marker expressed in activated B and plasma cells was positive in all cases, whereas CD138 staining was found in only a very small proportion of cells. Most samples contained diffuse CD3+ cells, but in all such cases these were a minor component. Flow cytometry analysis revealed that these CD3+ cells were B lymphocytes with aberrant expression of CD3, rather than a population of passenger T lymphocytes (Figure 3). Overall, these findings are consistent with post-transplant B cell lymphoma with the phenotype of non-germinal centre B cells.

**Figure 3:**
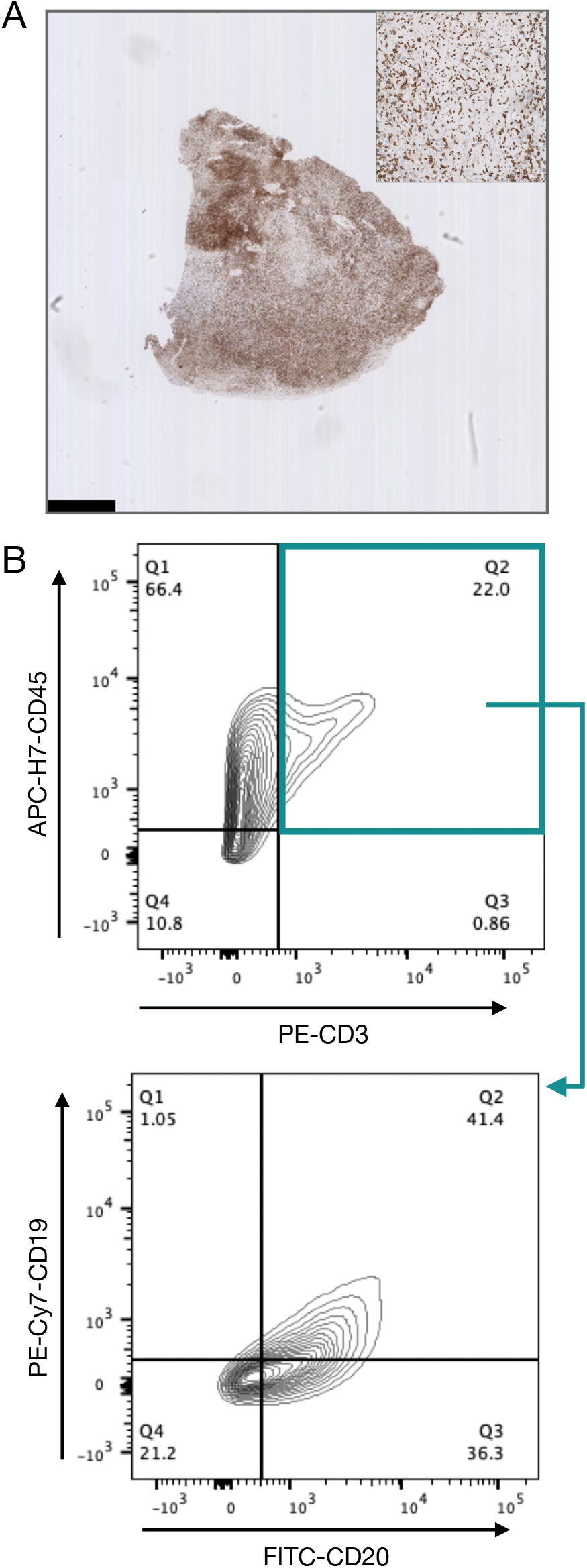
Inves3ga3on of CD3-expressing cells in B lymphoprolifera3ons. A) Immunohistochemical staining with an anF-CD3 anFbody in the CRUK0825 R7. Scale bar = 1 mm. Inset image is 500 μm in width. B) Flow cytometry analysis of CD19 and CD20 expression (lower panel) within CD45+/CD3+ cells (upper panel).

As EBV infection occurs in immunocompromised patients and transplant recipients^23^, we performed *in situ* hybridization for EBV-encoded small RNAs (EBER) to assess the EBV status of these lymphoproliferations. All lymphoproliferations were positive for EBER (15/15; Table 1; Supplementary File 1), which are expressed ubiquitously by EBV-infected cells regardless of their latency status^24^. It has not been possible to determine the EBV serology status of patients within the lung TRACERx cohort but none of the patients whose tumours generated lymphoproliferations in our study had a history of immunodeficiency, of lymphoproliferative disease or of prior therapy that would be expected to result in an immunodeficient state. Nor did we find evidence of germline mutations in any of these patients that would confer susceptibility to diffuse large B cell lymphoma (based on ^22^; data not shown).

**Table 1:**
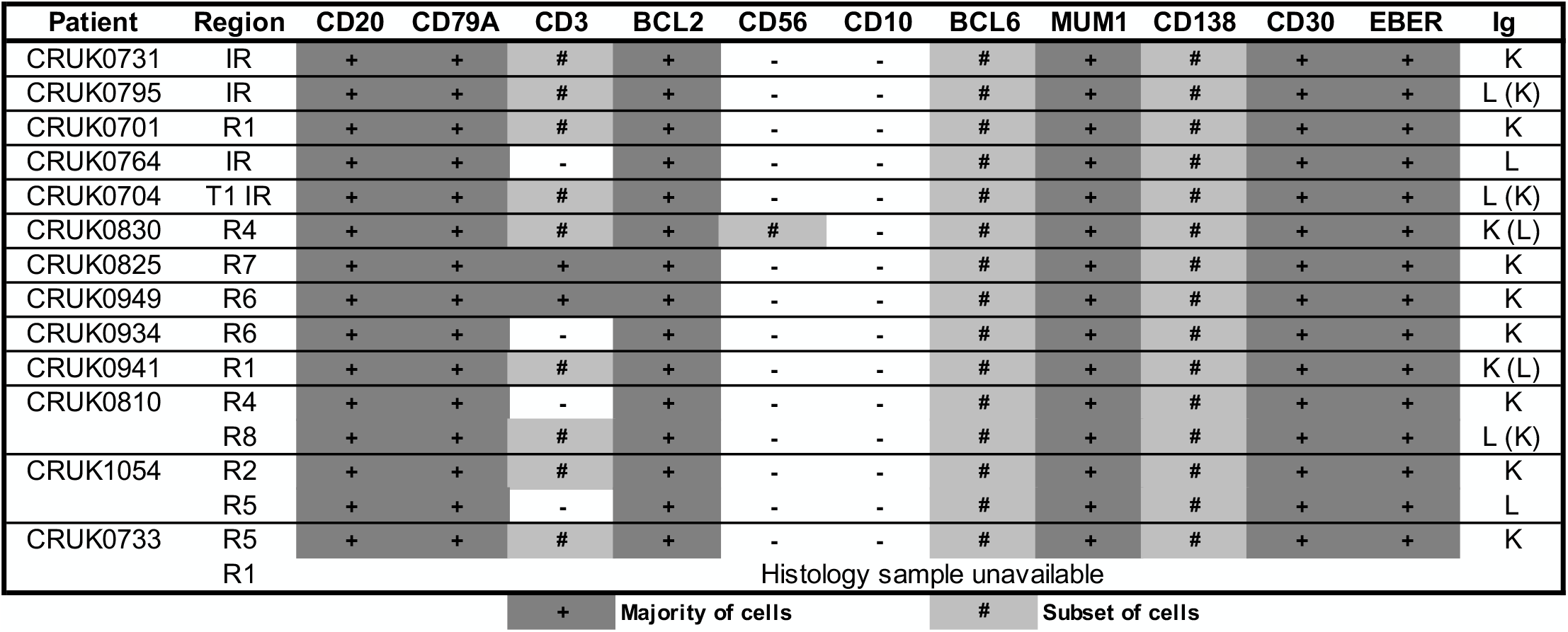
Immunophenotyping of B lymphoprolifera3ons arising in a NSCLC xenograD program. IR = inter-region (i.e. not matched to a tumour region defined within the lung TRACERx study). DetecFon of both IgK and IgL light chain expression by immunohistochemistry is indicated with the minor light chain shown in brackets.

### IgK and IgL rearrangements suggest independent origins of NSCLC PDX lymphoproliferations

Detection of immunoglobulin light chain restriction is indicative in routine diagnostic pathway for B-cell lymphoma. To evaluale the immunoglobulin light chain expression pattern in our samples, we performed single immunohistochemistry for kappa and lambda light chains. We found that 8/15 lymphoproliferations showed kappa only immunoglobulin (Ig) light chain restriction, two showing lambda light chain restriction and in the remaining five, co-expression of both kappa (IgK) and lambda (IgL) light chains was observed (Table 1; Figure 4A; Supplementary File 1). Dual expression of IgK and IgL has been shown in disease states ^25^ and at low frequency in healthy blood^26^. The presence of a cell population expressing both both IgK and IgL was confirmed by flow cytometry in two lymphoproliferations (CRUK0810 R8 and CRUK0941 R1) which had shown IHC evidence of dual expression (Figure 4B). For one xenograft (CRUK0733 R1), fixed tissue was unavailable for histological characterisation of IgK and IgL. We therefore used the standardised EuroClonality (BIOMED-2) PCR-based assay^27^ to determine the immunoglobulin light chain gene rearrangements in this sample and the two cases previously validated using flow cytometry. This assay uses pooled primers targeting rearrangements in the kappa locus (IGK, Figure 4C, left lane), rearrangements to the kappa deleting element (Kde), which occur once kappa rearrangements have been exhausted^28^ (Kde, middle lane), and subsequent rearrangements of the lambda locus (IGL, right lane). The neoplastic B cell nature of the CRUK0733 R1 xenograft was confirmed by the presence of two IGK rearrangements (Figure 4C).

**Figure 4:**
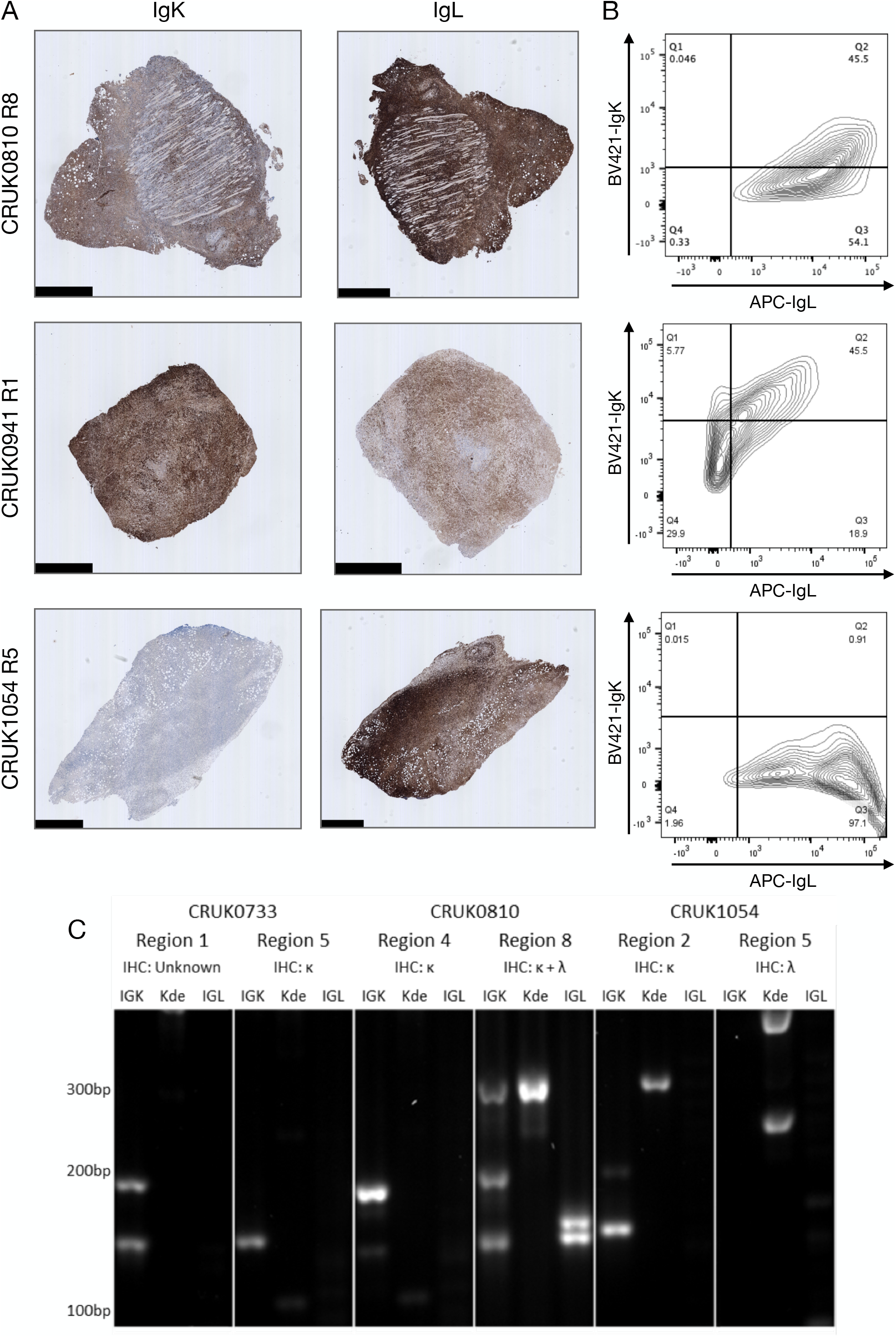
B lymphoprolifera3ons arising from different primary tumour regions show dis3nct immunoglobulin light chain rearrangements. A) Immunohistochemical staining with anF-IgK (le6) and anF-IgL (right) anFbodies in the CRUK0810 R8, CRUK0941 R1 and CRUK1054 R5 xenogra6s. Scale bars = 1 mm. B) Flow cytometry analysis of IgK and IgL expression within CD45+ cells that were also posiFve for either CD19 or CD20. C) PCR analysis of rearrangements to immunoglobulin light chain loci in mulF-region paFents including those to the kappa deleFng element (Kde).

Histological characterisation showed that lymphoproliferations arising from different spatial regions of the same primary tumour shared similar morphology and immunophenotype in all three cases for which multiple lymphoproliferations were available. However, distinct Ig light chain restrictions in lymphoproliferations arising from the same tumour indicated the likely independent origins of the lymphoproliferative xenografts in two cases for which immunohistochemistry was possible (Table 1; Figure 4; Supplementary File 1). In all three cases, PCR for Ig light chain gene rearrangements PCR demonstrated distinct rearrangements between the xenograft pairs, again supporting their independent clonal origins (Figure 4C).

Kappa and lambda light chain expression was detected in the CRUK0810 R8 xenograft by histology (Figure 4A) and flow cytometry (Figure 4B). The xenograft contained multiple rearrangements in IGK, Kde and IGL in the PCR assay (Figure 4C), suggesting the possibility that this lymphoproliferation may have contained multiple B cell clones. The CRUK1054 R5 xenograft demonstrated lambda light chain restriction by immunohistochemistry (Figure 4A) and flow cytometry (Figure 4B), but IGL rearrangement was not detected by PCR (Figure 4C). This xenograft did, however, show two rearrangements to Kde (Figure 4C), supporting progression to IGL rearrangement. The failure to detect IgL may have been caused by subsequent somatic hypermutation of IGL or by the use of an IGL gene segment not covered by the BIOMED-2 primer set^27^.

## DISCUSSION

In xenograft studies from a diverse range of tumour types, proliferations of non-tumour lymphocytes have been observed, a caveat in the development of pre-clinical cancer models. The rate of lymphoproliferation emergence varies between reports, with as many as 80% of growing lesions after three months being lymphoproliferations in one prostate cancer PDX pipeline^13,27^. It is feasible that age might be a factor in lymphoproliferation frequency in PDX pipelines but they have also been observed in paediatric solid cancers^29^. In lung cancer PDX experiments, lymphoproliferation rates of 16.7% in NOG mice (4/24;^17^), 12.4% in NOD-scid mice (19/153;^15^) and 11.5% in NSG mice (16/139; lung squamous cell carcinoma;^16^) have previously been observed. In accordance with these studies, we found a lymphoproliferation rate of 11.9% of all patient tissue injections. We did not observe more frequent lymphoproliferation formation in adenocarcinomas than squamous cell carcinomas as has been previously reported in a study using NOD-scid mice^15^. Over 25% of xenografts that formed in our study were lymphoproliferations and there was no difference in the time between implantation and harvest of lymphoproliferations and NSCLC PDXs that could be used to distinguish lymphoproliferations from true NSCLC PDX models. This indicates the relevance of screening PDX models using immunohistochemistry for keratin and CD45 expression as a rapid and relatively low-cost approach to identify lymphoproliferations in xenograft platforms. Screening models at the earliest opportunity and routinely during PDX passaging avoids the time-and cost-inefficient expansion of lymphoproliferations and mistaken tumour identity in downstream studies.

The origin of lymphoproliferations in the xenotransplantation setting is not fully understood. Given that tumour-infiltrating B cells are present in tumours, lymphoproliferations could feasibly arise from the activation and expansion of these cells. However, the association of lymphoproliferations with the reactivation of Epstein-Barr virus (EBV) – a herpes virus which persists as a life-long, asymptomatic latent infection in more than 90% of the human population and exhibits tropism for B lymphocytes^30^ – suggests that rare transformed cells that were previously kept in check by immune surveillance in patients can generate lymphoproliferations in immunocompromised mice. In immunocompetent patients, EBV infection would be expected to be confined to post-germinal centre memory cells^31^, it is therefore likely that these cells are present in the tumour tissue and will have undergone differentiation and immunoglobulin maturation. Indeed, the immunophenotype of the lymphoproliferations presented here is similar to that of post germinal centre cells and post-transplantation diffuse large B cell lymphoma. While the limited analysis of Ig light chain amplicon length by agarose gel electrophoresis presented here is not sufficient to determine if a single lymphoproliferation is monoclonal, it does demonstrate that lymphoproliferations arising from independent spatial regions of primary tumours were clonally distinct. These data suggest that tumours contain multiple clones of EBV-transformed memory B cells possessing proliferative potential in NSG mice, where immunosurveillance by patient T cells is absent.

One approach to preventing the formation of lymphoproliferations might therefore be to quantify EBV RNA in starting material to filter out regions most likely to contain these B cells. Our finding that cryopreservation has little impact on NSCLC PDX take rate while not selectively favouring either tumour or B cell proliferations, would allow a time window in which to implement this approach^19^. However, a previous study in NSCLC has suggested that the extent of EBER positivity in tissue did not predict lymphoproliferation formation^15^. Monoclonal antibody therapy against CD20 using rituximab results in B cell depletion and is an approved therapy in some leukaemias and B cell non-Hodgkin lymphoma. Injection of mice with a single dose of rituximab at the point of xenograft implantation has been shown to reduce lymphoproliferation formation in ovarian^14^ and hepatobiliary/gastrointestinal PDX platforms. However, tumour histology affected the efficacy of this approach^25^, so further data are required on its efficacy in NSCLC PDX models given the substantial cost of dosing every first passage mouse in large PDX derivation studies.

In conclusion, post-transplant-like diffuse large B cell proliferations were a frequent outcome in our NSCLC xenograft study using NSG mice. Here we present the characterisation of these lymphoproliferations, including several cases in which lymphoproliferations were derived from spatially distinct regions of the primary tumour. In these cases, we demonstrate that the proliferations likely arise from distinct cells of origin. None of the patients from which these lymphoproliferations were derived suffered haematological malignancy diagnosis with 3.5 years of follow up. This suggests that there are multiple B cell clones with lymphoproliferative potential within a primary NSCLC tumour and that there is continuous surveillance of these cells within the tumour. Our data highlight the value of quality control measures to identify lymphoproliferations within xenograft pipelines and support the incorporation of strategies to either minimise lymphoproliferations during the early stages of xenograft establishment pipelines.

## METHODS

### Generation and maintenance of xenograft models

Ethical approval to generate patient-derived models was obtained through the Tracking Cancer Evolution through Therapy (TRACERx) clinical study (REC reference: 13/LO/1546; https://clinicaltrials.gov/ct2/show/NCT01888601). Animal studies were approved by the University College London Biological Services Ethical Review Committee and licensed under UK Home Office regulations (P36565407).

Tissue from patients undergoing surgical resection of non-small cell lung cancers was immediately transported on ice from theatres to a pathology laboratory where it was dissected for diagnostic and then research purposes. Region-specific tumour samples were dissected by a consultant pathologist such that the tissue used to generate patient-derived xenograft (PDX) models was spatially adjacent to the tissue that was sequenced in TRACERx. Individual region-specific tumour samples were transported to the laboratory in transport medium consisting of MEM alpha medium (Gibco) containing 1X penicillin/streptomycin (Gibco), 1X gentamicin (Gibco) and 1X amphotericin B (Fisher Scientific, UK), minced using a scalpel and resuspended in 200 µl growth factor-reduced Matrigel (BD Biosciences). Tumour tissue was minced with a scalpel rather than dissociated to single cell suspensions to preserve local cytoarchitecture. To generate PDX tumors, male non-obese diabetic/severe combined immunodeficient (NOD/SCID/IL2Rγ^-/-^; NSG) mice were anaesthetized using 2–4% isoflurane, the right flank was shaved and cleaned before minced tumour tissue in matrigel was injected subcutaneously using a 16G needle. Mice were observed during recovery, then regularly monitored for tumour growth. Mice were kept in individually ventilated cages under specific pathogen-free conditions and had *ad libitum* access to sterile food and autoclaved water. Tumour monitoring was performed twice per week and tumour measurements taken in two dimensions using callipers. When xenograft tumours formed, mice were terminated before tumours reached 1.5 cm^3^ in volume. Mice without xenograft tumours were terminated at a median of 306 days (range 37-402 days). Successfully engrafted tumours were propagated by injection of minced xenograft tissue in matrigel into a new host, with banking of FFPE tissue, OCT-embedded tissue and DNA at each generation. Cryopreservation of patient material and xenograft tissue at each tumour transfer was performed in foetal bovine serum plus 10% DMSO.

### Immunohistochemical characterisation

Formalin-fixed paraffin-embedded (FFPE) tissue sections were routinely obtained at PDX passage by fixation of tumour fragments (approximately 3×3×3 mm in size) in 4% paraformaldehyde overnight at 4°C. Samples were stored in 70% ethanol at 4°C before being processed and embedded in paraffin. FFPE tissue sections of PDX tumours and their equivalent primary tumour region were subjected to hematoxylin and eosin (H&E) staining or immunohistochemistry using the following antibodies: anti-human CD45 (Clone: HI30; 1:200; BioLegend, #304002), anti-keratin (Clone: AE1/AE3; 1:100; Agilent #13160), anti-CD3 (Clone: LN10; 1:100; Leica Biosystems, #NCL-L-CD3-565), anti-CD20 (Clone: L26; 1:200; Agilent, #M0755), anti-CD30 (Clone: JCM182; RTU; Leica Biosystems, #PA0790), anti-CD56 (Clone: 564; RTU; Leica Biosystems, PA0191), anti-CD79A (Clone: JCB117; 1:100; Agilent; #M7050), anti-CD138 (Clone: MI15; 1:100; Agilent; #M7228), anti-BCL2 (Clone: BCL-2/100/D5; RTU; Leica Biosystems; #PA0117), anti-BCL6 (Clone:LN22; RTU; Leica Biosystems; #PA0204), anti-MUM1 (Clone:MUM1p; 1/400; Agilent; #M7259), anti-IgL (Clone: N/A; 1:400; Agilent; #GA507), and anti-IgK (Clone: N/A; 1:4000; Agilent; #A0191). Optimization of the antibodies and staining conditions was carried out on sections of human tonsil. Immunostaining was performed using the automated BOND-III Autostainer (Leica Microsystems, UK) according to protocols described previously^32^. Slide images were acquired using a NanoZoomer 2.0HT whole slide imaging system (Hamamatsu Photonics, Japan). Supplementary File 1 showing semi-automatically generated overview images with a selected region of interest was created from Nanozoomer ndpi whole-slide digital images by a custom python module using the openslide package^33^. This tool (PATHOverview) is available on Github (https://github.com/EpiCENTR-Lab/PATHOverview).

### EBER in situ hybridisation

*In situ* hybridization for detection of the EBV-encoded small RNAs (EBER) was performed using the EBER Probe (Catalogue no: ISH5687; Leica Biosystems Newcastle Ltd, UK). The technique was carried out on an automated stainer, BOND-III Leica Biosystems) platform according to the manufacturer’s instructions.

### PCR of Ig light chain

DNA was extracted from xenografts using either the PureLink Genomic DNA Mini Kit (Invitrogen) or the DNA/RNA AllPrep Kit (Qiagen). PCR for IGK, Kde and IGL along with a positive control ladder reaction was performed on 35-120 ng DNA utilising the BIOMED-2^27^ protocol with pooled primers detailed in Supplementary Table 1. ABI Gold Buffer (Applied Biosystems) was used in a 25 µl reaction. IGK, Kde and positive control reactions were performed with 1.5 mM MgCl_2_ and IGL with 2.5mM MgCl_2_. Cycling conditions were: 95°C for 7 min followed by 35 cycles of 95°C for 30s, 60°C for 30s, 72°C for 30s with a final extension at 72°C for 10min. PCR products were imaged on a 1% agarose gel.

### Flow cytometry

Cryopreserved xenografts were dissociated by mashing through a 30 μm cell strainer and blocked using 10% foetal bovine serum in PBS. Cells were resuspended in staining buffer consisting of 50% PBS containing 1% bovine serum albumin (BSA) and 50% Brilliant Stain Buffer (BD Biosciences) and incubated for 20 minutes at 4°C in the following antibodies: anti-CD45 (APC-H7), anti-CD3 (PE), anti-CD19 (PE-Cy7, Thermo Fisher Scientific, 25-0199-41), anti-CD20 (FITC; Biolegend, 302303), anti-IgK (BV421; Biolegend, 392705) and anti-IgL (APC; Biolegend, 316609). Cells were washed and re-suspended in flow cytometry buffer (PBS + 1% BSA) for flow cytometry. Cells were analysed using a BD LSRFortessa X-20 Cell Analyser (UCL Division of Medicine Core Facility, University College London) and data were analysed in FlowJo (v10).

### Resource availability

Biological materials, including xenograft models generated within this study, will be made available to the community for academic non-commercial research purposes via standard MTA agreements. PATHOverview, the Python-based tool used to generate histology overview images in this manuscript, has been made available via GitHub (https://github.com/EpiCENTR-Lab/PATHOverview).

## ACKNOWLEDGEMENTS

The authors thank the members of the TRACERx consortium for their contributions to this study. TRACERx (Clinicaltrials.gov no: NCT01888601) is sponsored by University College London (UCL/12/0279) and was approved by an independent research ethics committee (REC 13/LO/1546). TRACERx is funded by Cancer Research UK (CRUK; C11496/A17786) and is coordinated by CRUK and the UCL Cancer Trials Centre. The authors thank Sharon Vanloo for administrative support. **T.K**. is supported by the Japan Society for the Promotion of Science (JSPS) overseas research fellowships program (202060447). **N.M**. is a Sir Henry Dale Fellow, jointly funded by the Wellcome Trust and the Royal Society (211179/Z/18/Z), and also receives funding from CRUK, the Rosetrees Trust, the NIHR BRC at University College London Hospitals and the CRUK University College London Experimental Cancer Medicine Centre. **V.R**. received funding support from a MRC-CARP fellowship (MR/T024968/1). This work was also supported by a Medical Research Council grant (MR/P00184X/1) to **A.N.A. C.S**. is a Royal Society Napier Research Professor (RSRP\R\210001). This work was supported by the Francis Crick Institute that receives its core funding from Cancer Research UK (CC2041), the UK Medical Research Council (CC2041), and the Wellcome Trust (CC2041). C.S. is funded by Cancer Research UK (TRACERx (C11496/A17786), PEACE (C416/A21999) and CRUK Cancer Immunotherapy Catalyst Network); Cancer Research UK Lung Cancer Centre of Excellence (C11496/A30025); the Rosetrees Trust, Butterfield and Stoneygate Trusts; NovoNordisk Foundation (ID16584); Royal Society Professorship Enhancement Award (RP/EA/180007); NIHR University College London Hospitals Biomedical Research Centre; the Cancer Research UK-University College London Centre; Experimental Cancer Medicine Centre; the Breast Cancer Research Foundation (US) BCRF-22-157; Cancer Research UK Early Detection and Diagnosis Primer Award (Grant EDDPMA-Nov21/100034); and The Mark Foundation for Cancer Research Aspire Award (Grant 21-029-ASP). This work was supported by a Stand Up To Cancer-LUNGevity-American Lung Association Lung Cancer Interception Dream Team Translational Research Grant (Grant Number: SU2C-AACR-DT23-17 to S.M. Dubinett and A.E. Spira). Stand Up To Cancer is a division of the Entertainment Industry Foundation. Research grants are administered by the American Association for Cancer Research, the Scientific Partner of SU2C. C.S. is in receipt of an ERC Advanced Grant (PROTEUS) from the European Research Council under the European Union’s Horizon 2020 research and innovation programme (grant agreement no. 835297). **R.E.H**. was supported by a Sir Henry Wellcome Postdoctoral Fellowship (Wellcome Trust; WT209199/Z/17) and received additional funding for this project from the CRUK Lung Cancer Centre of Excellence, the Roy Castle Lung Cancer Foundation and the James Tudor Foundation. R.E.H. is a National Institute for Health and Care Research (NIHR) Great Ormond Street Hospital (GOSH) Biomedical Research Centre (BRC) Collaborative Catalyst Fellow. The views expressed are those of the author(s) and not necessarily those of the NHS, the NIHR or the Department of Health. For the purpose of open access, the author has applied a CC BY public copyright licence to any Author Accepted Manuscript version arising from this submission.

