## Supplementary File 1 for "Phenotyping of lymphoproliferative tumours generated in xenografts of non-small cell lung cancer"

CRUK0731_IR_P0

IHC outcome: Lymphoma. Patient diagnosis: Squamous cell carcinoma.

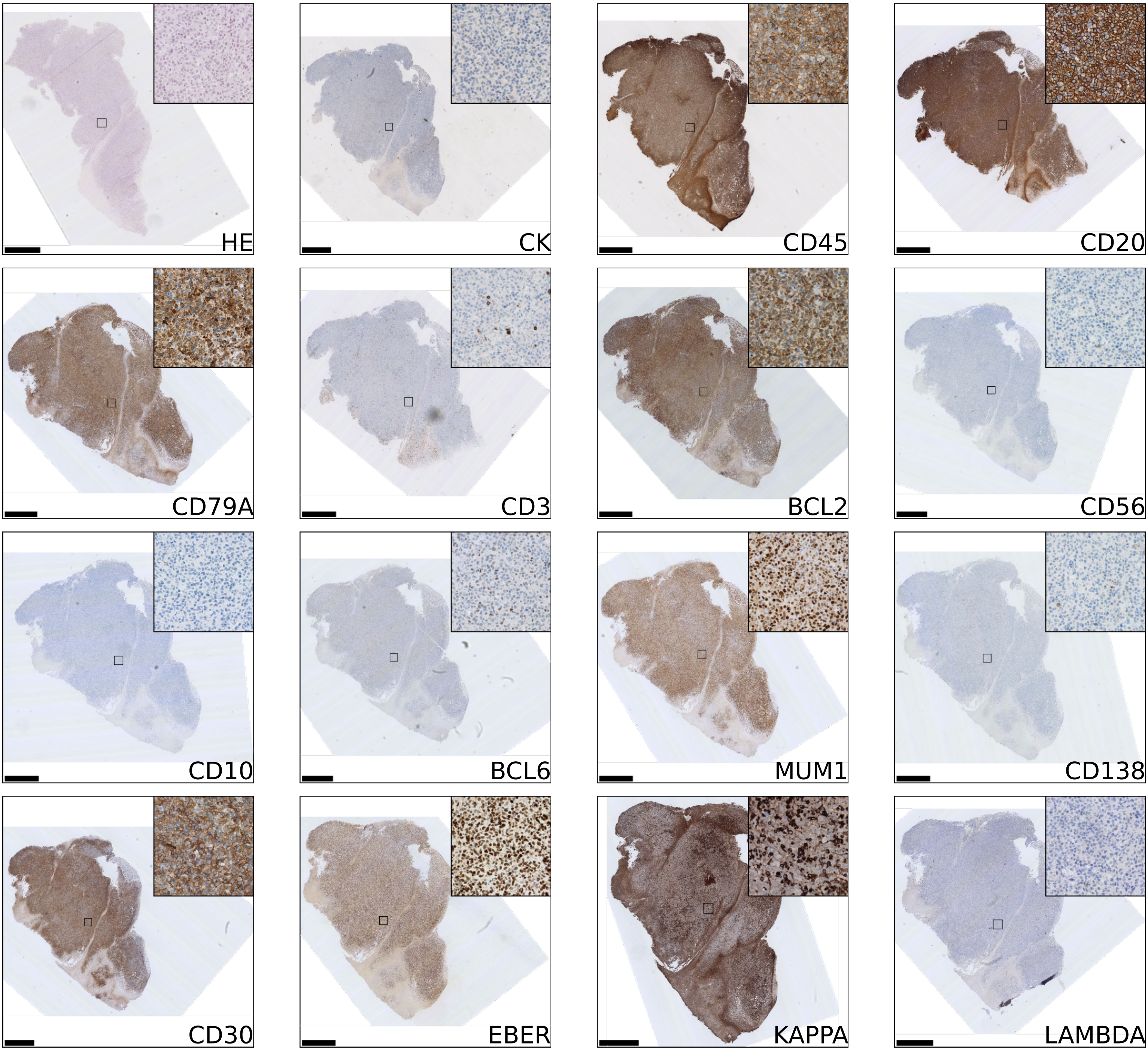


Scale bar 1.0mm. Inset image width 250um.

CRUK0795_IR_P0

IHC outcome: Lymphoma.    Patient diagnosis: Invasive adenocarcinoma.

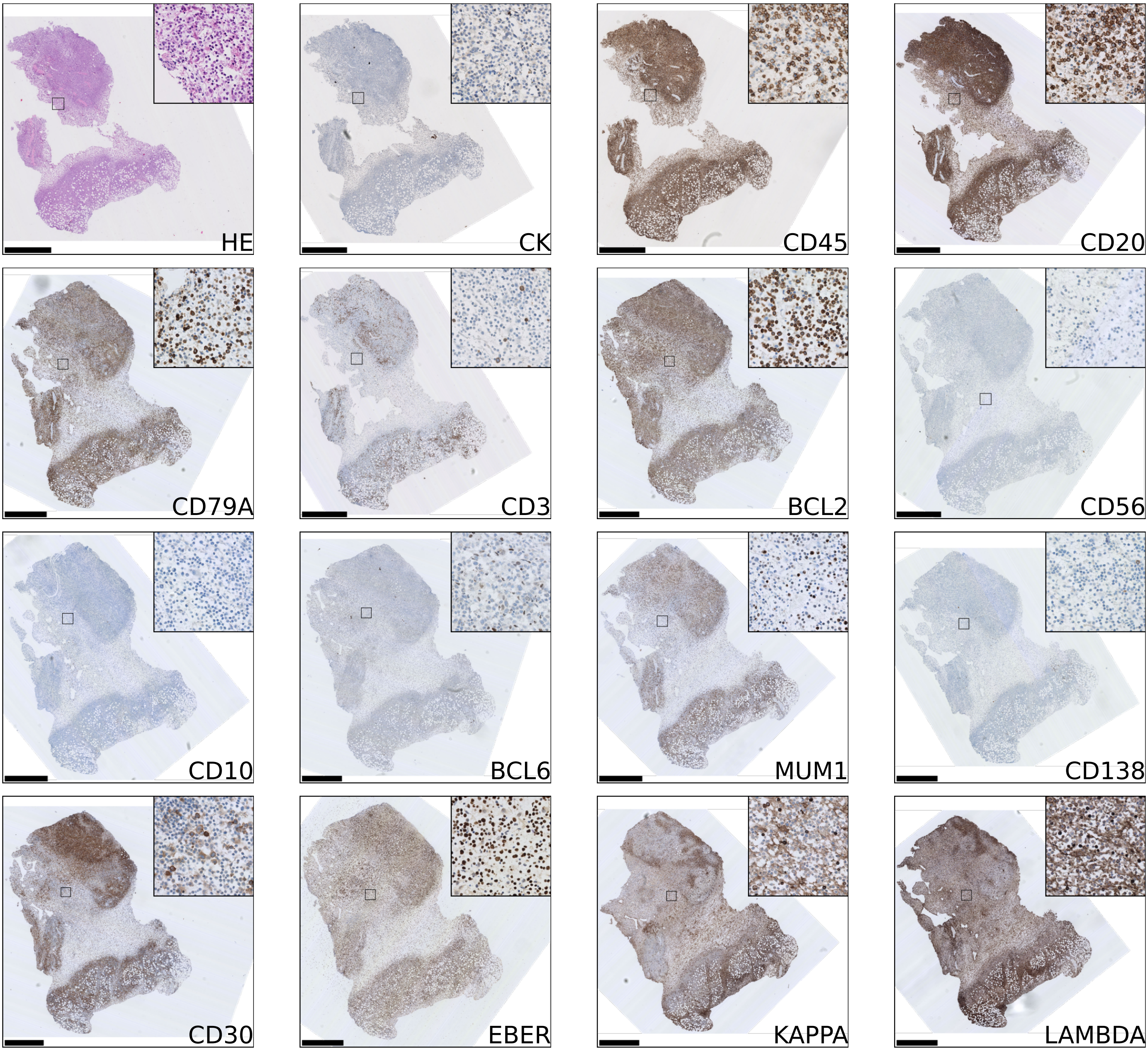


Scale bar 1.0mm. Inset image width 250um.

CRUK0701_R1_P0

IHC outcome: Lymphoma.   Patient diagnosis: Squamous cell carcinoma.

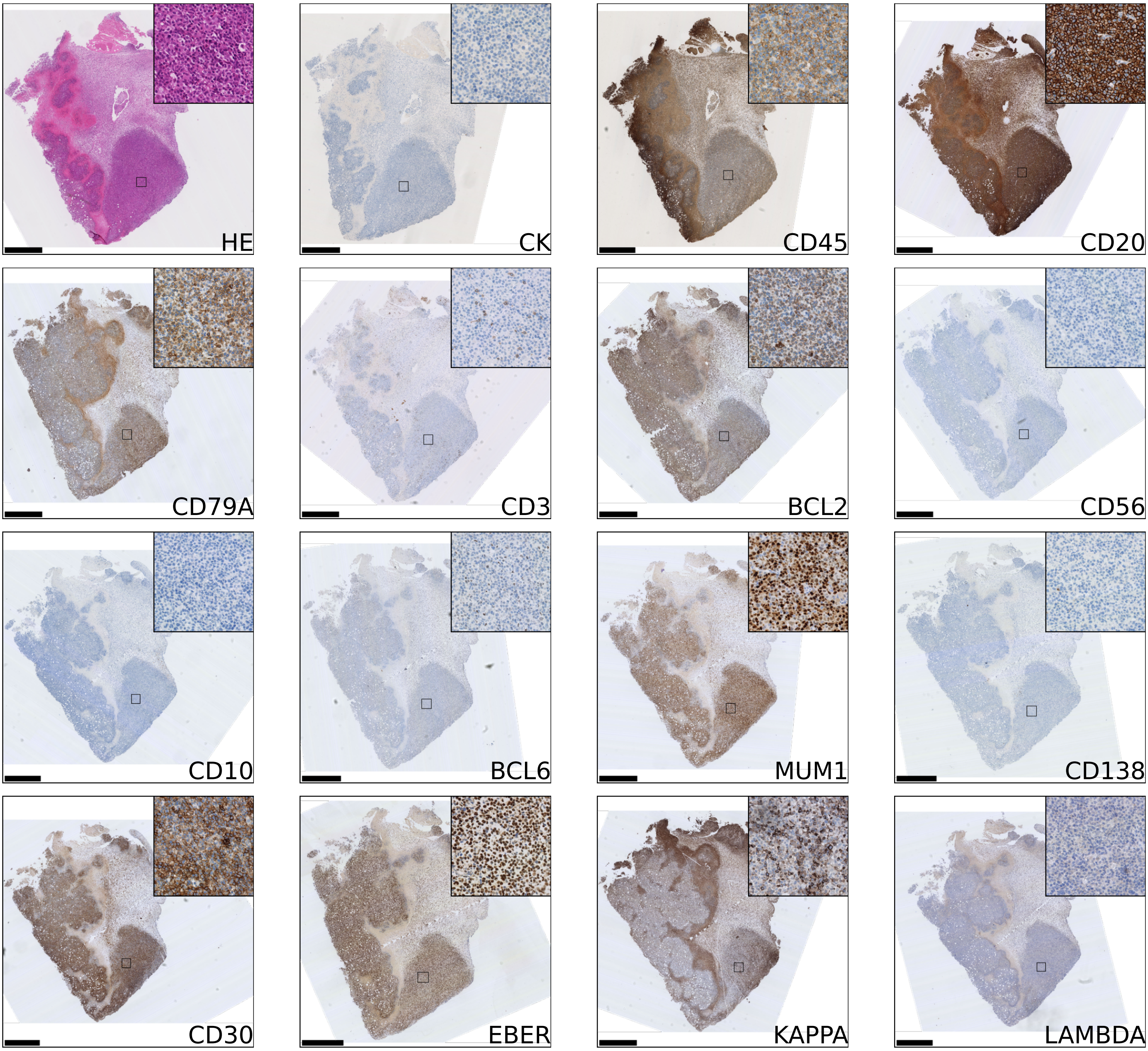


Scale bar 1.0mm. Inset image width 250um.

CRUK0733_R5_P0

IHC outcome: Lymphoma. Patient diagnosis: Invasive adenocarcinoma.

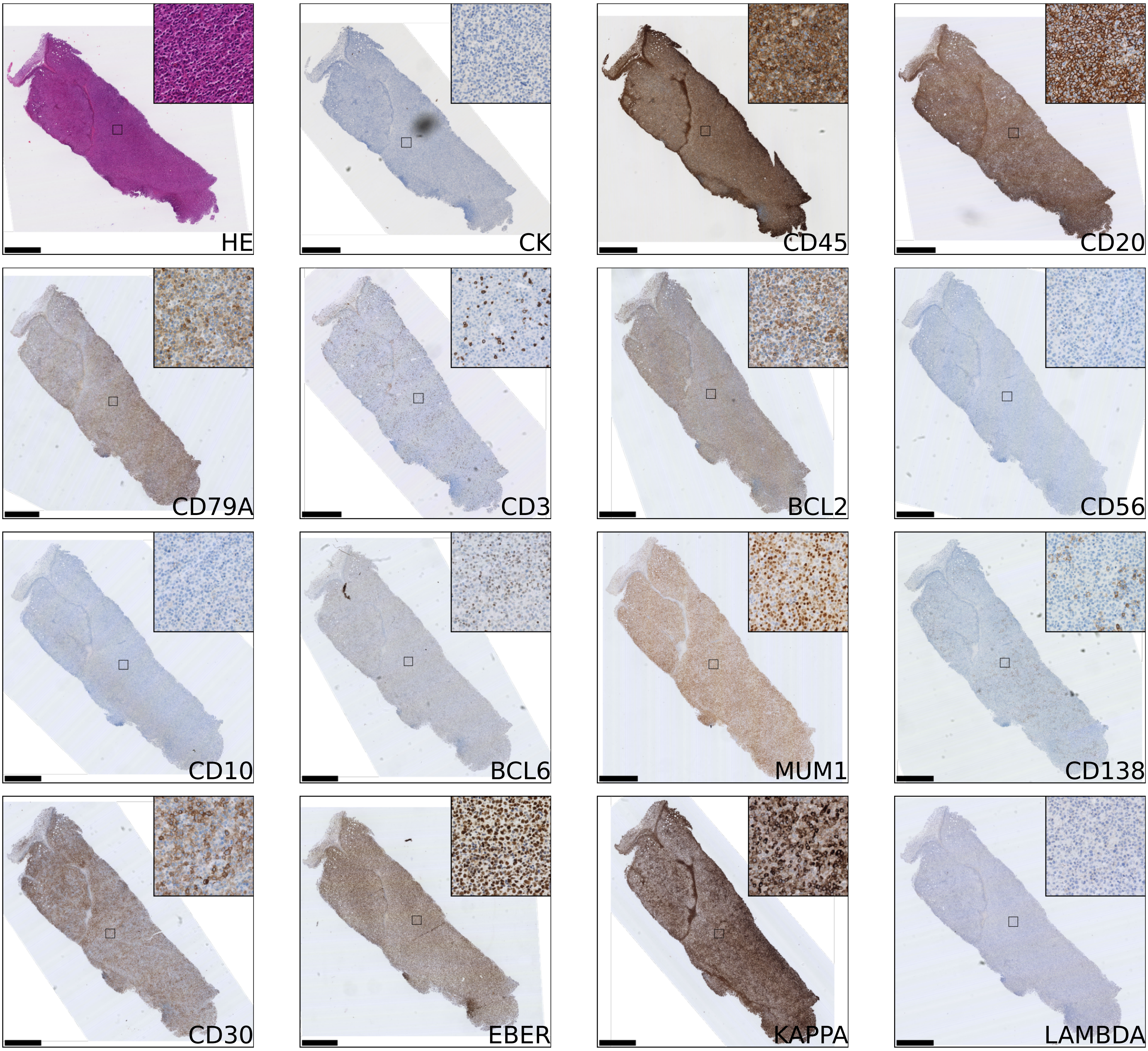


Scale bar 1.0mm. Inset image width 250um.

CRUK0764_IR_P0

IHC outcome: Lymphoma.    Patient diagnosis: Invasive adenocarcinoma.

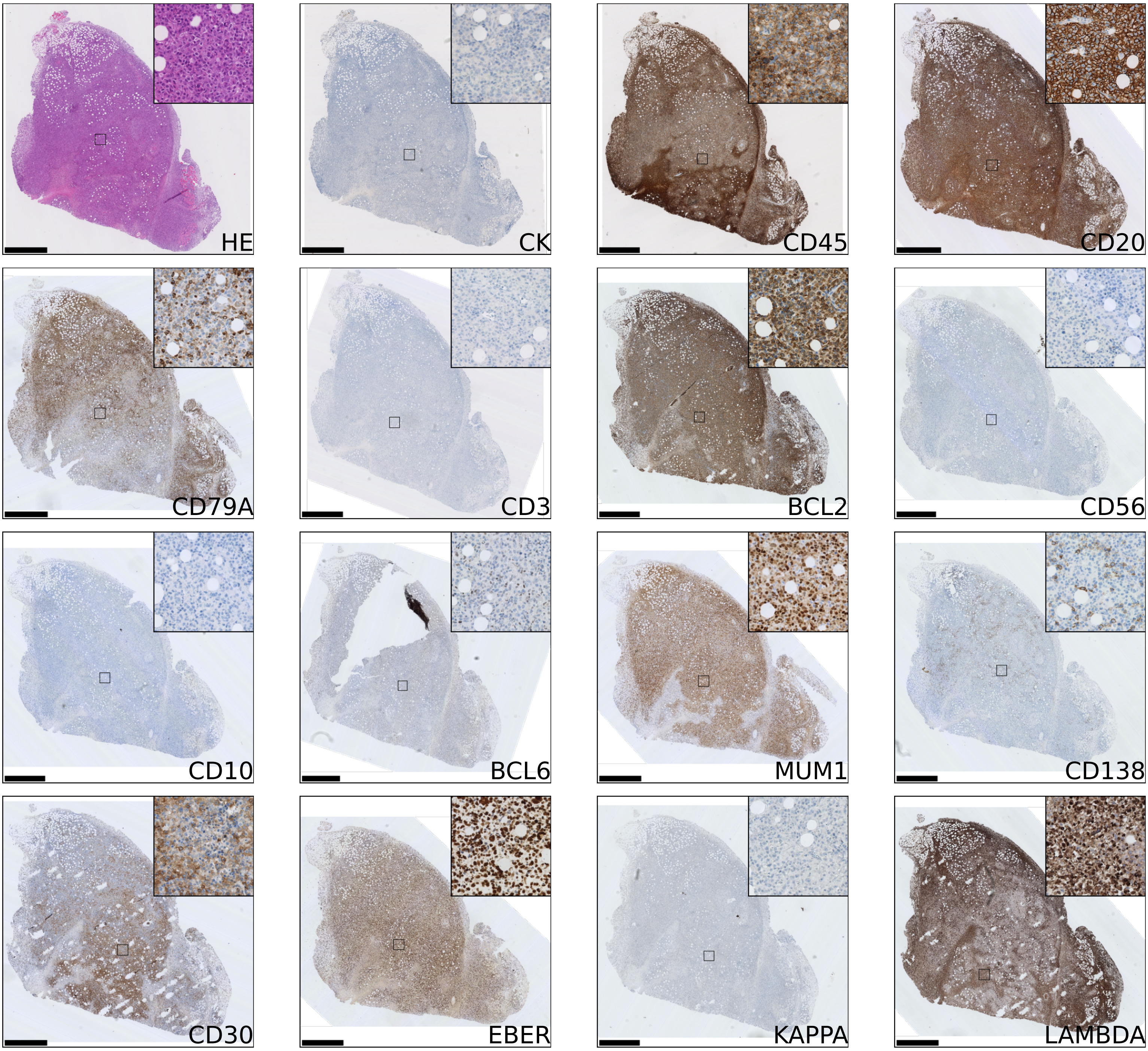


Scale bar 1.0mm. Inset image width 250um.

CRUK0704_T1-IR_P0

IHC outcome: Lymphoma.   Patient diagnosis: Invasive adenocarcinoma.

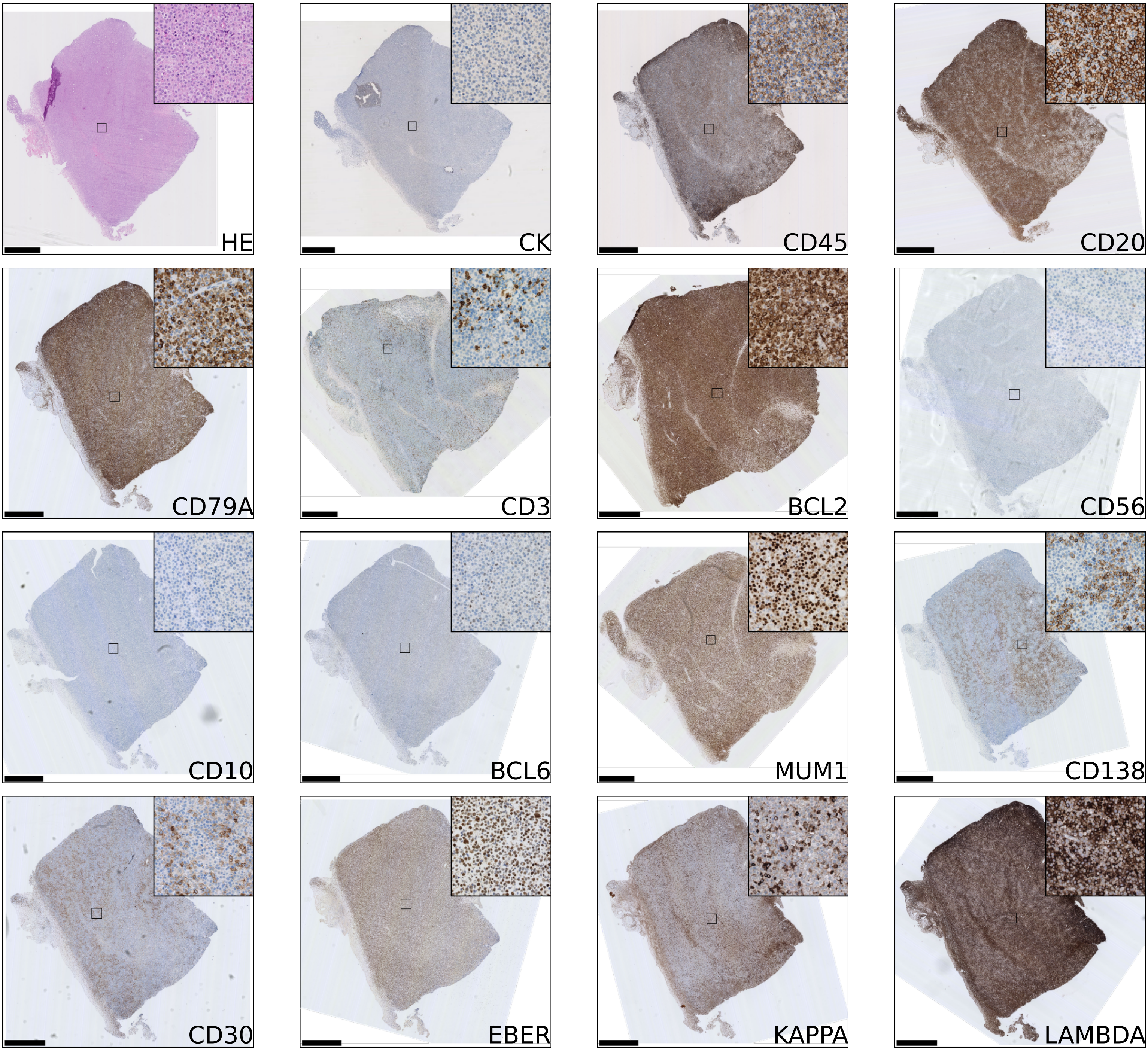


Scale bar 1.0mm. Inset image width 250um.

CRUK0810_R4_P0

IHC outcome: Lymphoma. Patient diagnosis: Invasive adenocarcinoma.

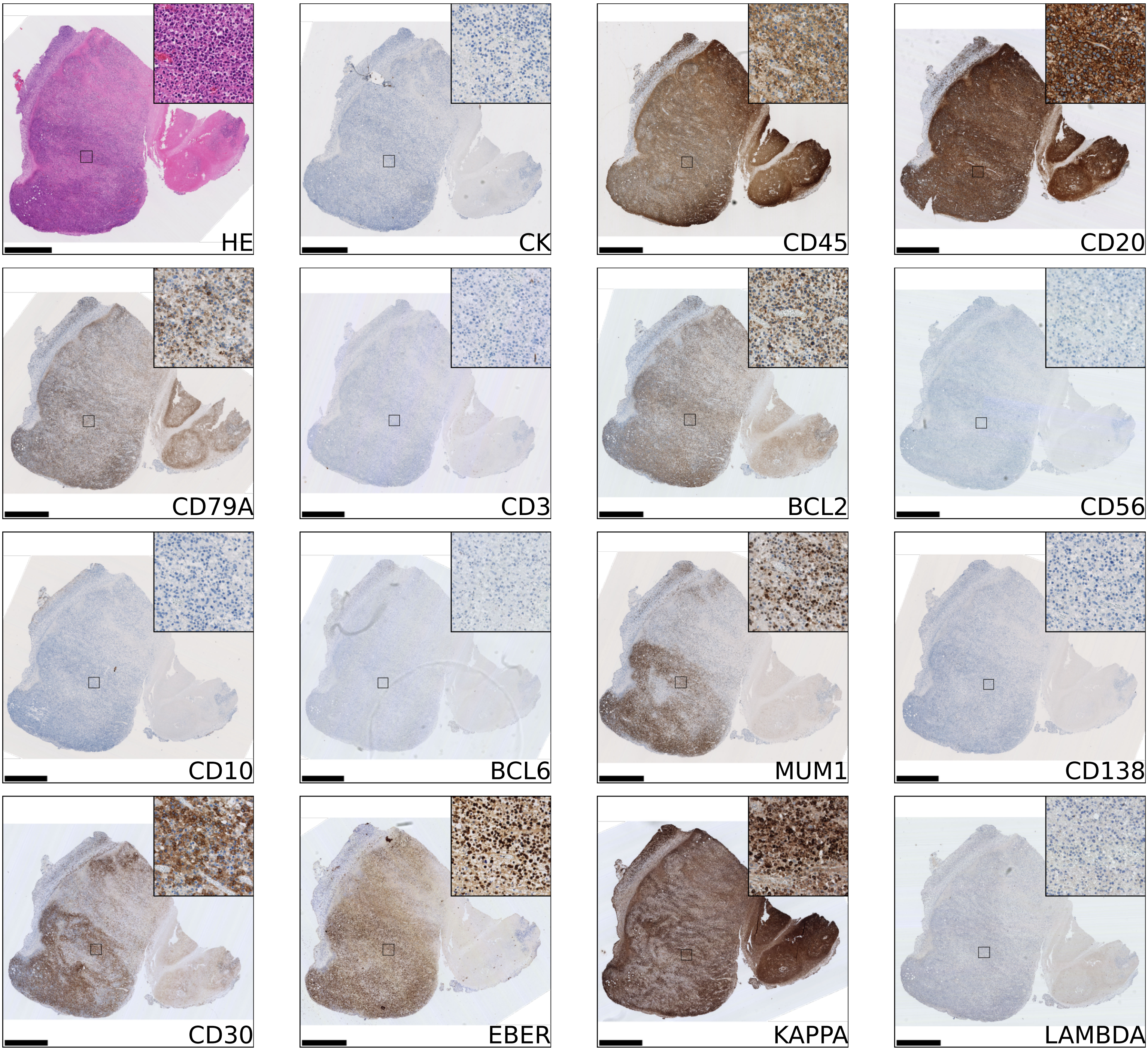


Scale bar 1.0mm. Inset image width 250um.

CRUK0810_R8_P0

IHC outcome: Lymphoma.   Patient diagnosis: Invasive adenocarcinoma.

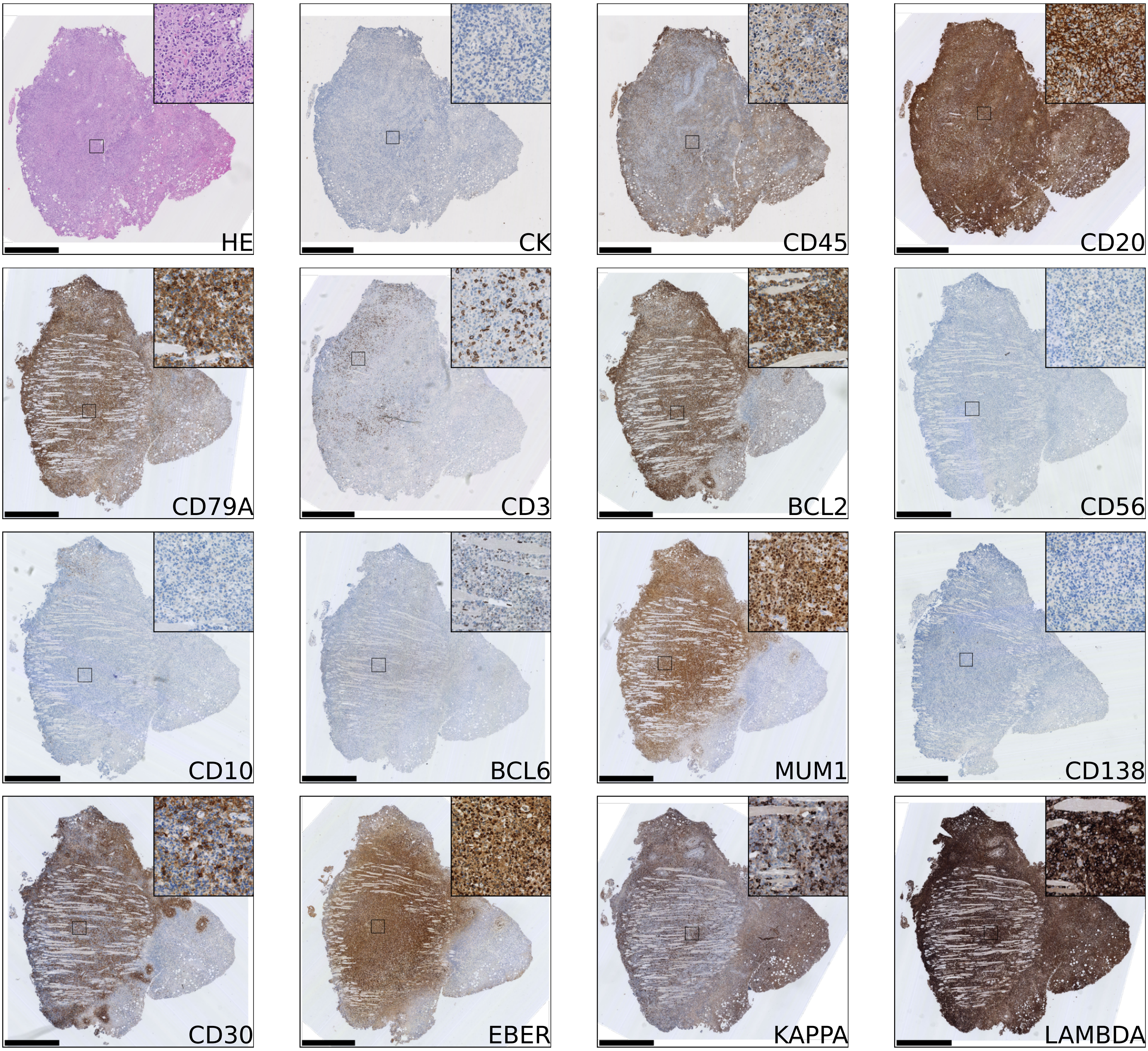


Scale bar 1.0mm. Inset image width 250um.

CRUK0830_R4_P0

IHC outcome: Lymphoma.   Patient diagnosis: Other.

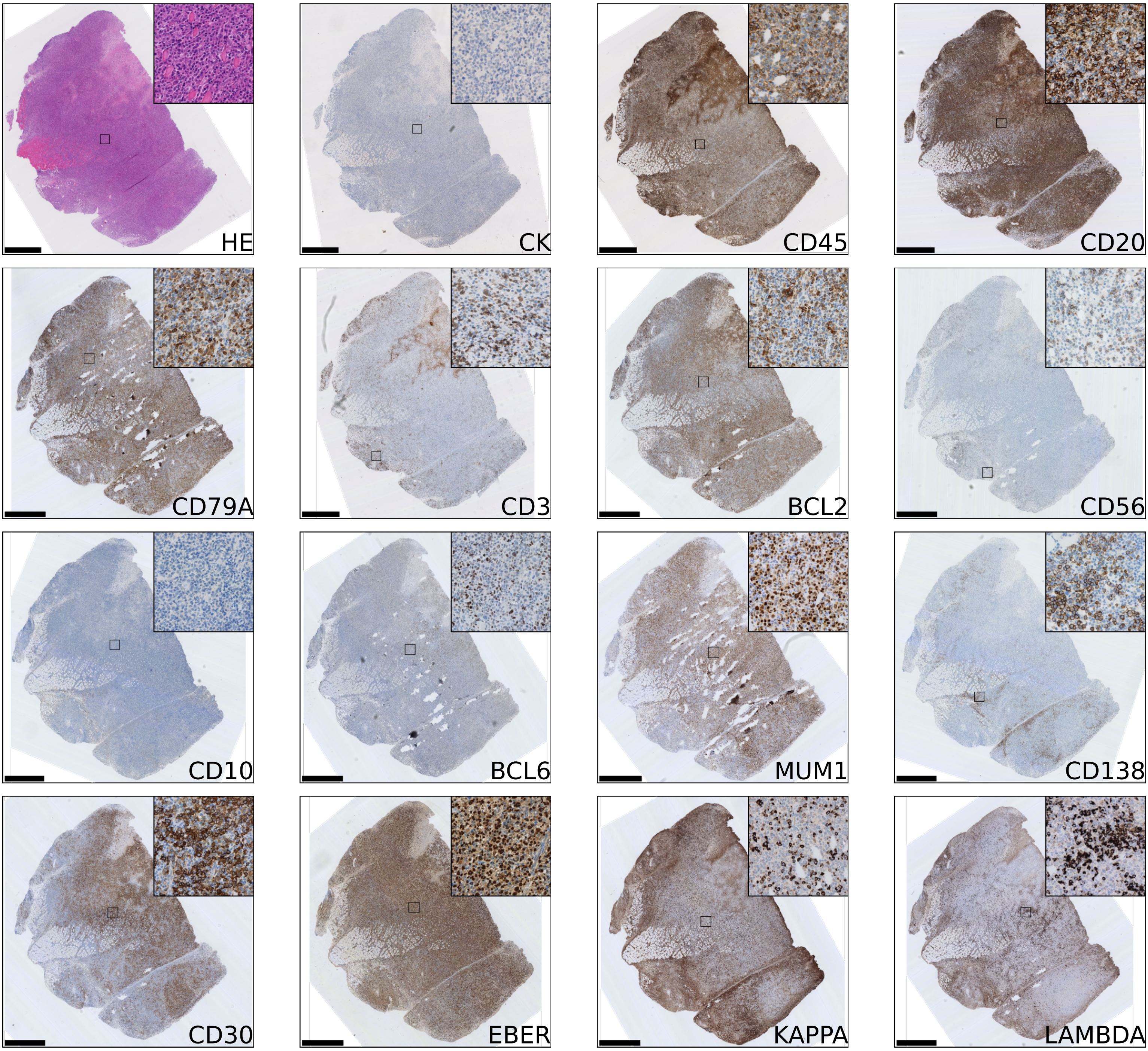


Scale bar 1.0mm. Inset image width 250um.

CRUK0825_R7_P0
IHC outcome: Lymphoma. Patient diagnosis: Invasive adenocarcinoma.

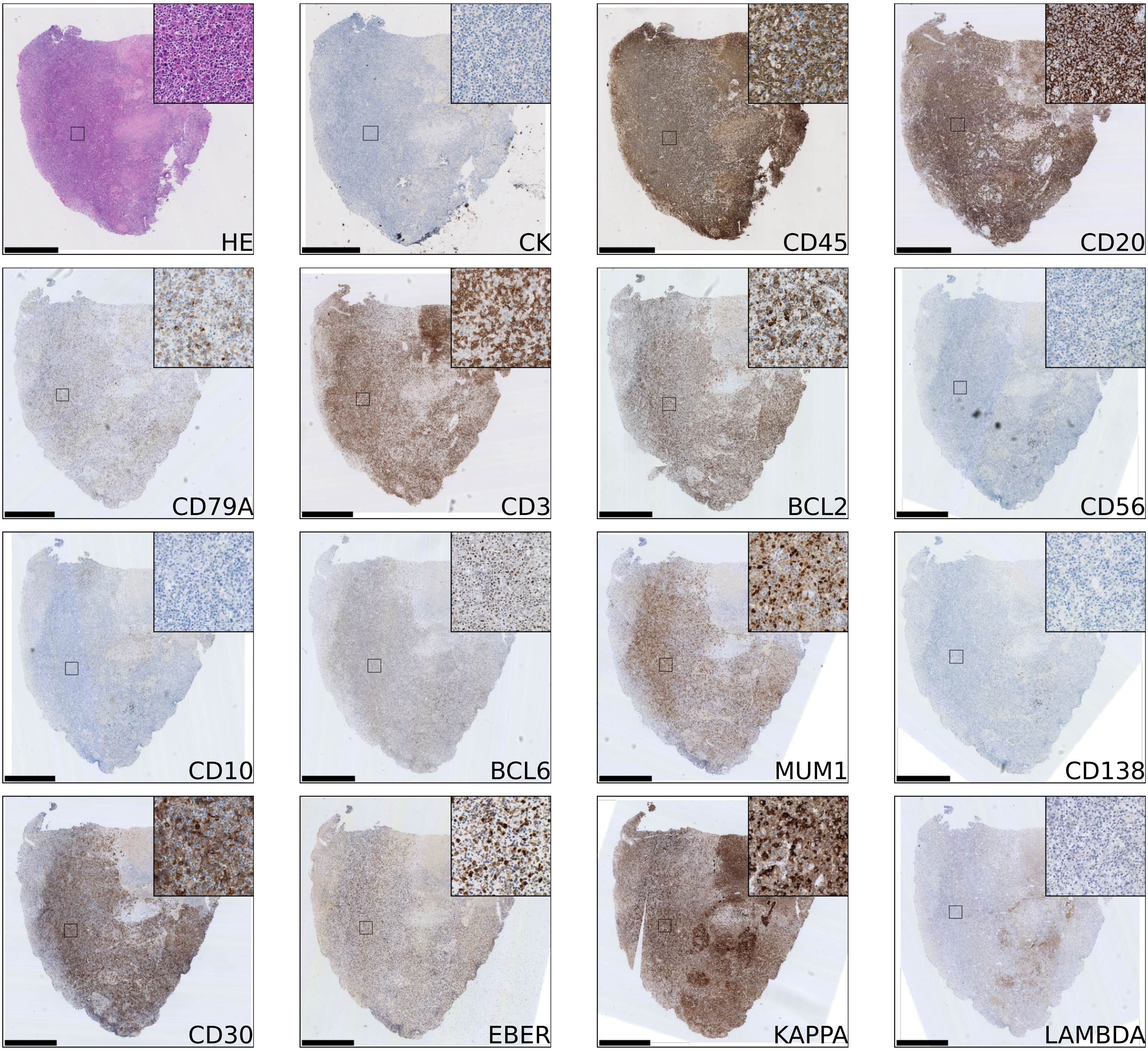


Scale bar 1.0mm. Inset image width 250um.

CRUK0949_R6_P0
IHC outcome: Lymphoma. Patient diagnosis: Squamous cell carcinoma.

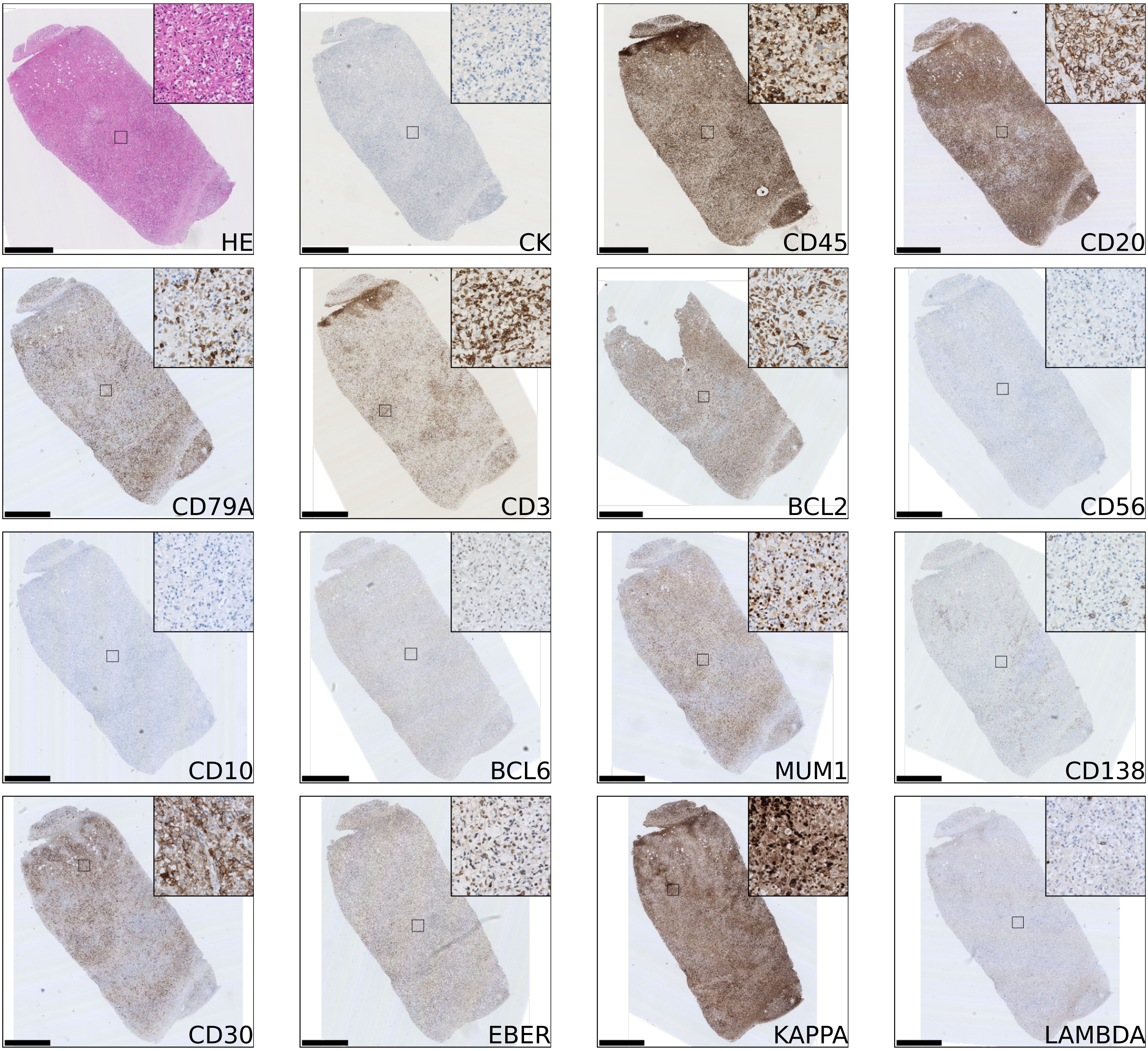


Scale bar 1.0mm. Inset image width 250um.

CRUK0934_R6_P0

IHC outcome: Lymphoma. Patient diagnosis: Squamous cell carcinoma.

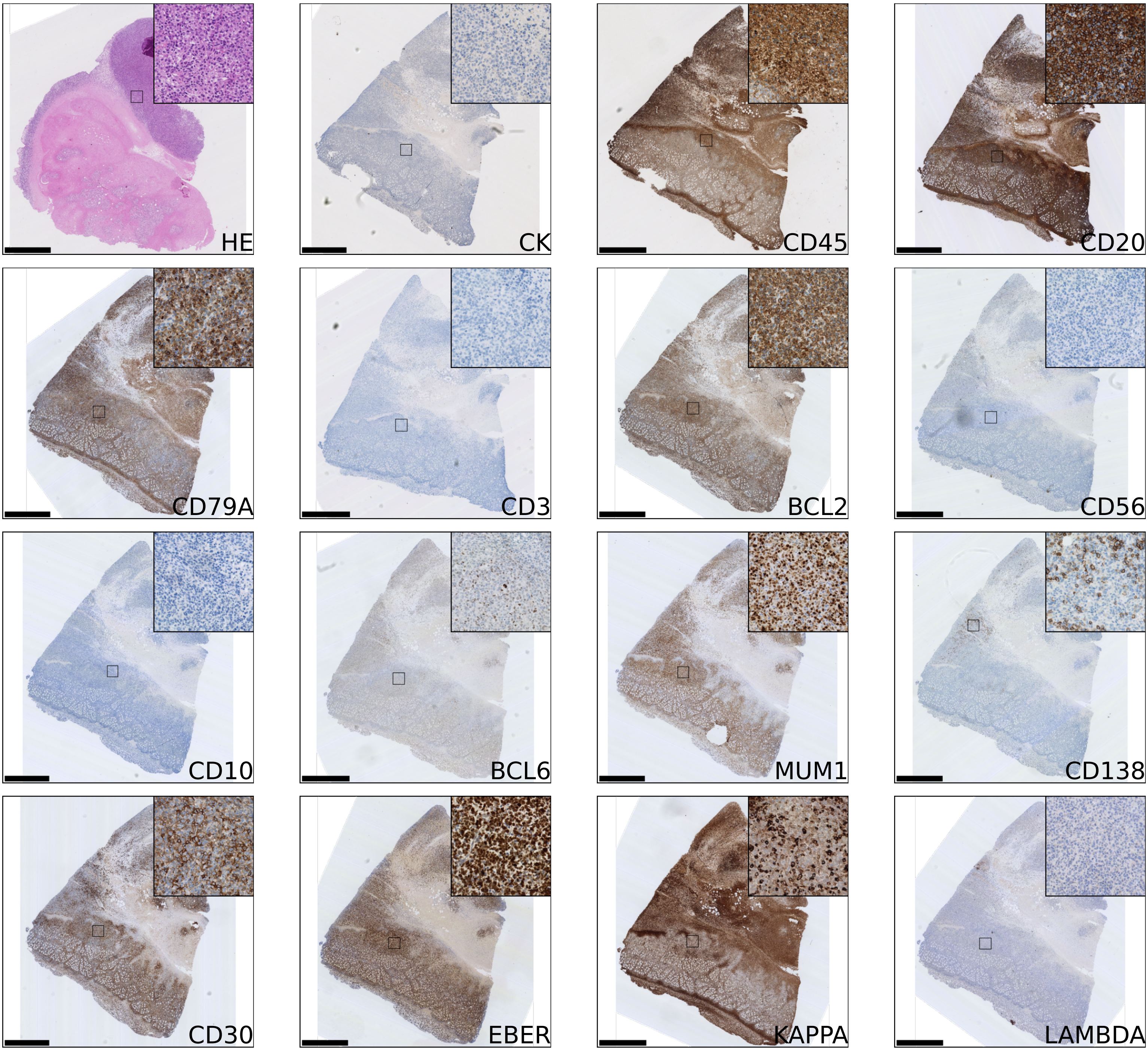


Scale bar 1.0mm. Inset image width 250um.

CRUK0941_R1_P0

IHC outcome: Lymphoma.   Patient diagnosis: Invasive adenocarcinoma.

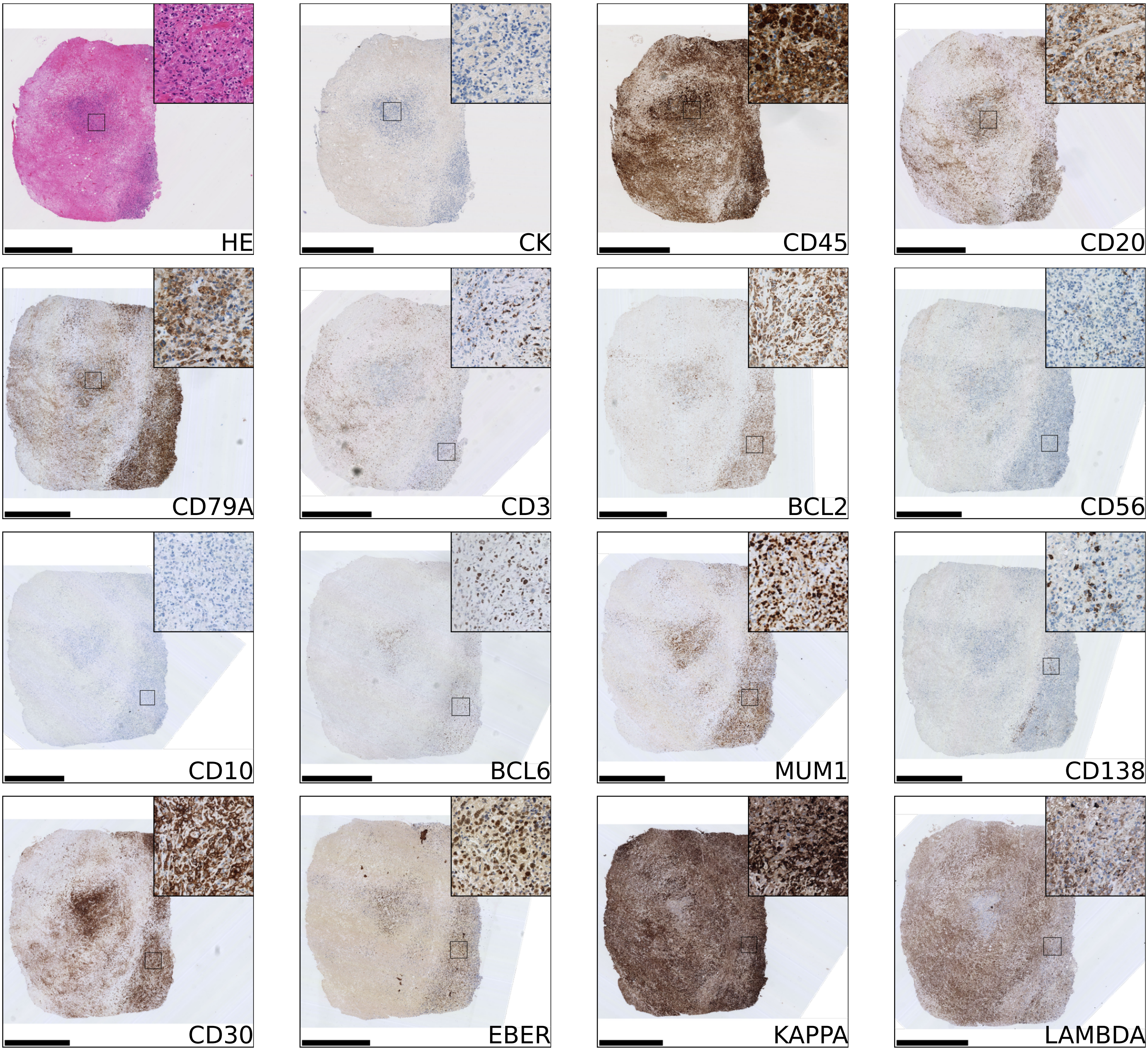


Scale bar 1.0mm. Inset image width 250um.

CRUK1054_R2_P0

IHC outcome: Lymphoma.   Patient diagnosis: Squamous cell carcinoma.

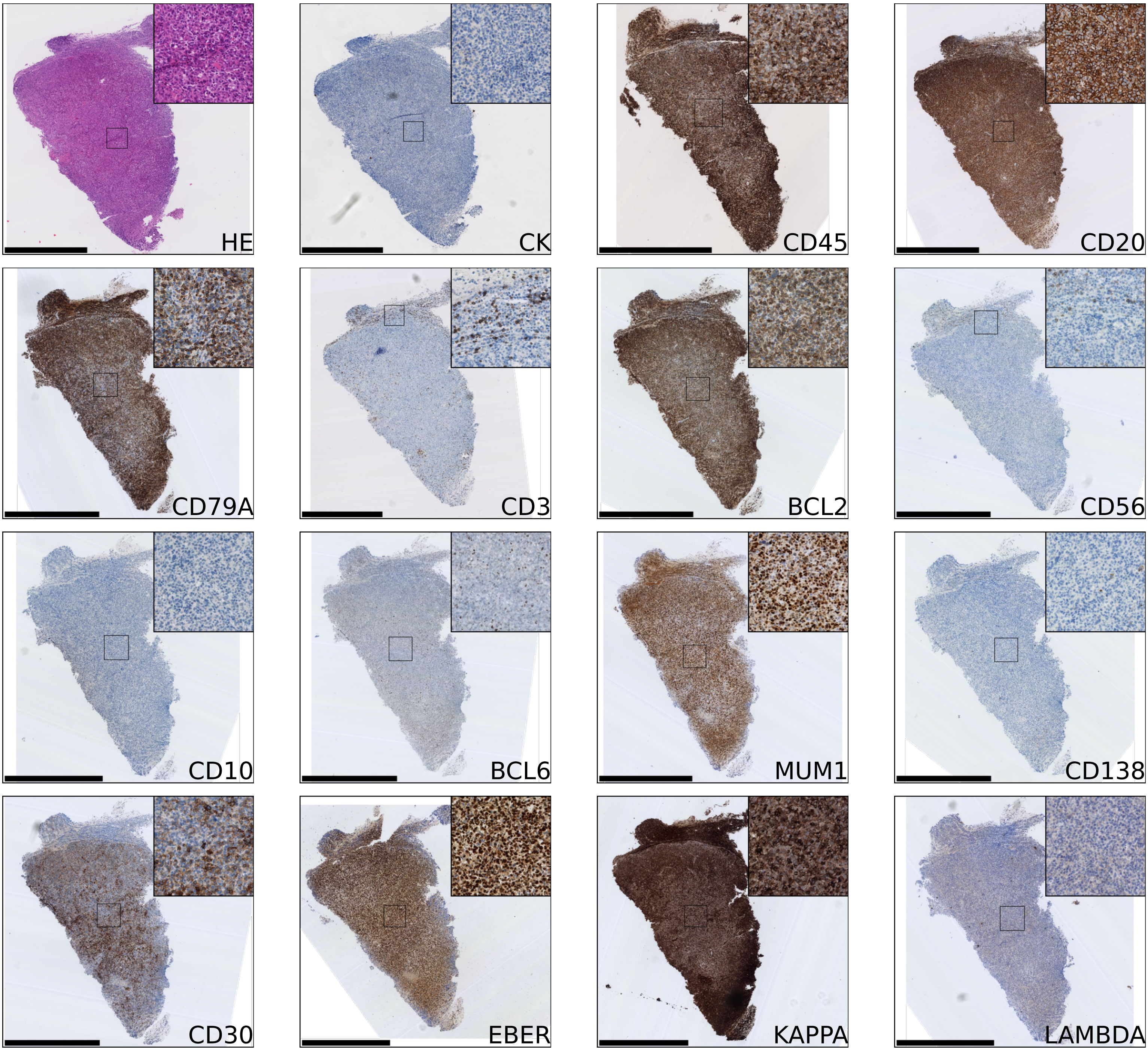


Scale bar 1.0mm. Inset image width 250um.

CRUK1054_R5_P0
IHC outcome: Lymphoma.   Patient diagnosis: Squamous cell carcinoma.

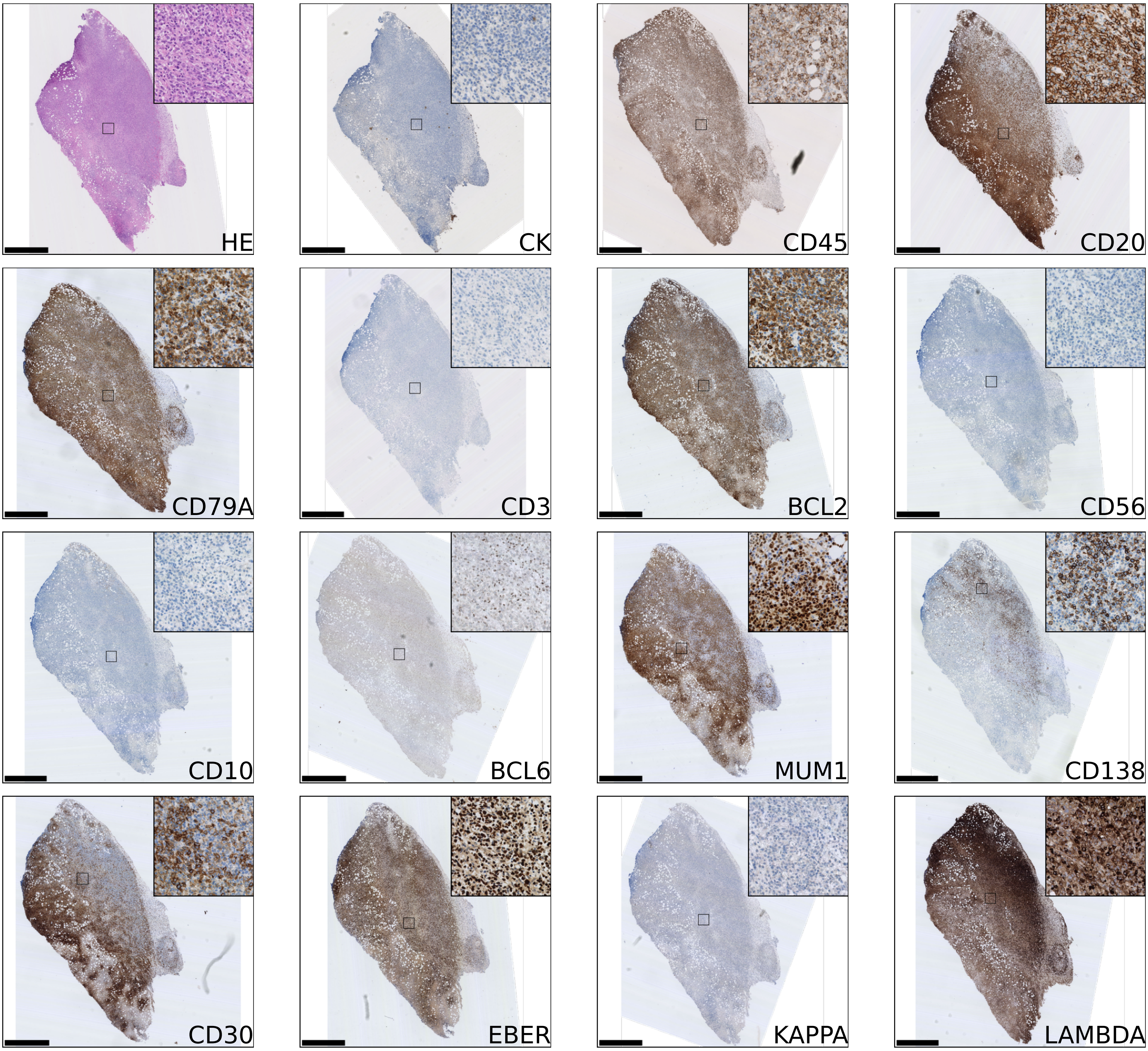


Scale bar 1.0mm. Inset image width 250um.
